# Salt-inducible kinase (SIK) inhibition is protective in a mouse model of asthma

**DOI:** 10.1101/2021.06.07.447230

**Authors:** Manuel van Gijsel-Bonnello, Nicola J. Darling, Laura McDonald, Mairi Sime, Jonathan Clark, Mokdad Mezna, Alexander Schuettelkopf, Craig Mackay, Justin Bower, Heather McKinnon, Philip Cohen, J. Simon C. Arthur

## Abstract

Interleukin-13 (IL-13) and other Th2 cytokines are important regulators of airway hyper-responsiveness, immune cell infiltration and inflammation in allergic asthma, and are produced when immune cells, such as type 2 innate lymphoid cells (ILC2s) and mast cells, are stimulated with IL-33. Here, we report that the IL-33-dependent secretion of IL-13 from ILC2s is prevented by inhibition of the salt-inducible kinases (SIKs), as we have shown previously in mast cells (Darling NJ *et al.*, 2021, J. Biol. Chem. doi: 10.1016/j.jbc.2021.100428). We also report that a new SIK inhibitor with improved pharmacokinetic properties suppresses the recruitment of eosinophils to the lungs and serum immunoglobulin E (IgE) levels in the *Alternaria alternata-induced* model of allergic asthma. Our results suggest that drugs targeting SIK isoforms may have therapeutic potential for the treatment of asthma.

## Introduction

It has been estimated that over 300 million people worldwide suffer from asthma, making it the most prevalent chronic respiratory disease [1]. Key mediators of asthma include the Th2 cytokines, such as interleukin-13 (IL-13) and IL-5, which are produced by several types of immune cells, including type 2 innate lymphoid cells (ILC2s) and mast cells [2]. These mediators drive some of the characteristic features of asthma, such as airway inflammation, smooth muscle constriction and mucus secretion [2, 3]. In chronic asthma, eosinophils are also recruited to the lungs [3–5] and asthma patients have a higher eosinophil count in both the lungs and peripheral blood [6]. Indeed, asthma severity and the frequency of exacerbations is correlated with eosinophil count in many (but not all) asthma subtypes [6, 7].

In humans, allergic sensitisation to fungal pathogens, such as *Alternaria alternata*, is a risk factor for asthma and correlates with asthma severity and the exacerbation to fatal asthma [8, 9]. In mice, administration of an extract of *Alternaria* induces allergic lung inflammation, including both the recruitment of eosinophils to the lungs and the production of immunoglobulin E (IgE), and is therefore used as a mouse model for allergic asthma. IgE provides an important link between the antigen recognition role of the adaptive immune system and the effector functions of mast cells, where it primes the IgE-mediated allergic response by binding to Fc receptors [10]. ILC2s may be important for the development of inflammation in the *Alternaria* model [11–14]. The model appears to be strongly influenced by IL-33 [15], since an IL-33 antagonist inhibits the inflammatory response [16]. IL-33 stimulates the production of a number of Th2 related cytokines, including IL-13, IL-5 and IL-4, which together promote a number of processes associated with asthma, including eosinophil recruitment and IgE production by plasma cells [2, 17].

IL-33 is released from airway epithelial cells following physical damage or necrosis [17]. It is upregulated in asthma patients, and the level of IL-33 correlates with disease severity [18, 19]. In addition, polymorphisms in human IL-33 or the IL-33 receptor subunit have been linked to the incidence of asthma [20]. IL-33 stimulates ILC2s and mast cells to secrete IL-13 and other cytokines, such as granulocyte-macrophage colony stimulating factor (GM-CSF) [21–25]. IL-13 promotes airway hyper-responsiveness, the overproduction of mucus and infiltration of eosinophils into the lungs, resulting in airway obstruction [26–28]. Airway administration of GM-CSF *in vivo*, alongside the house dust mite model of asthma, lowers the threshold for IL-13 and IL-5 secretion and eosinophil recruitment to the lungs. Administration of GM-CSF in this model increases the expression of IL-33 in lung epithelial cells and the resultant cytokine secretion and eosinophil recruitment is IL-33-dependent [29]. GM-CSF also promotes the proliferation and maturation of antigen-presenting cells, which are critical for the presentation of allergens to T cells [30].

We have recently reported that the IL-33-stimulated transcription of the genes encoding IL-13 and GM-CSF and the secretion of these cytokines from mast cells is suppressed by small molecule inhibitors of all three SIK isoforms. IL-13 and GM-CSF secretion was also suppressed in mast cells expressing kinase-inactive mutants of both SIK2 and SIK3 or lacking any expression of these two SIKs [23]. Here, we show that the IL-33-dependent secretion of IL-13 and other cytokines is also suppressed by SIK inhibition in ILC2s and that the secretion of GM-CSF is reduced partially in ILC2s. These findings suggested that SIK inhibitors might be protective in mouse models of asthma. In this study we use a new SIK inhibitor that is highly selective with good pharmacokinetic properties to demonstrate that SIK inhibition suppresses eosinophil recruitment and IgE production in the *Alternaria alternata*-induced model of allergic asthma. Our results suggest that drugs targeting SIK2 and SIK3 may have therapeutic potential for the treatment of asthma.

## Materials and Methods

### Materials

Recombinant IL-33, IL-2 and IL-7 were purchased from Peprotech, reconstituted in sterile water, and stored in aliquots at −80 °C.

### Compound B

Compound B is a potent, and selective bioavailable SIK inhibitor suitable for oral dosing in mice. Previous work has found that compound B has excellent affinity for SIK2 as measured in a biochemical IMAP assay using full length SIK protein, with modest selectivity over SIK1 and SIK3, and >100 fold selectivity over a wider panel of 137 kinases (Invitrogen). Using a cellular translocation assay (DiscoverX), Compound B was shown to have <20 nM potency in cells, and 100% bioavailability following oral dosing in CD1 female mice with a T_max_ at 4 h. Blood samples were collected from mice at termination, placed in EDTA tubes to prevent coagulation and mixed 1:1 in distilled water before storage at −20°C or processing. Proteins were precipitated with acetonitrile and compound concentrations measured by UHPLC-TOF mass spectrometry using electrospray ionisation, in comparison to calibration standards. The pharmacokinetic profile for Compound B in mice following a single oral gavage dose of 10 mg/kg in 25 % (v/v) propylene glycol/1 % (v/v) Tween/PBS vehicle, is shown in **Fig S1**.

### Mast cell culture, purity and cytokine secretion

Bone marrow-derived mast cells (BMMC) were generated from the bone marrow of mice aged 3-6 months and their purity was analysed as described previously [23]. Cytokine concentrations in cell free culture medium or serum samples were measured using the Bio-Plex Pro Assay System (Bio-Rad Laboratories) [23]. Serum IgE concentrations were measured using an ELISA kit (Biolegend) according to the manufacturer’s instructions.

### Type 2 innate lymphoid cell culture

ILC2s were isolated from the mesenteric fat of wild type (WT) mice aged 3-6 months old, using 4-6 mice per culture as described previously [25]. The antibodies used are detailed in [25], with dilutions and cell washes performed using ice-cold PBS containing 0.5 % (w/v) BSA and 2 mM EDTA. Lineage negative, CD45 positive cells were collected, and up to 10^5^ cells were plated in 0.2 ml of cell culture medium (RPMI-1640, 10 % (v/v) heat-inactivated FBS, 100 U/ml penicillin, 100 μg/ml streptomycin, 4 mM L-glutamine, 10 mM HEPES, 1 mM sodium pyruvate, 50 μM ß-mercaptoethanol, 20 ng/ml IL-2, 10 ng/ml IL-7) and incubated for 5 days at 37 °C with 5 % CO_2_. On the second day, half of the culture medium was removed and replaced with fresh medium without disturbing the cells. ILC2s were seeded in 96 well plates at 3000-5000 cells per well in 100 μl media without IL-2 and IL-7, and rested for 1 h prior to experimentation.

### Mice

C57Bl6/J mice were obtained from Charles River Laboratories (UK) and were allowed a minimum of 15 days acclimatisation following shipment before any experimental interventions. Animals were maintained under specific pathogen-free conditions consistent with E.U. and U.K. regulations. Mice were housed in individually ventilated cages at 21 °C, 45-55 % humidity, and a 12/12 h light/dark cycle, with free access to food (RM3) and water. This work was performed under a U.K. Home Office Project Licence awarded after recommendation by the University of Dundee Ethical Review Committee.

### *Alternaria*-induced allergic lung inflammation

Age matched female 7-8 week old C57Bl6/J mice were acclimatised to rehydrated powdered food (powdered RM3, Special Diet Services, Witham, Essex) twice during the week before the start of the experiment (in the presence of normal diet) to avoid neophobia. Compound B was administered in the food to avoid stress that may be caused by gavage dosing over the 35 day time course of the experiment. Two days prior to the start of compound addition to the food, mice were moved exclusively to the rehydrated powdered food. Compound B was dissolved at 20 mg/ml in DMSO and kept in the dark at −20 °C until required. Compound B or vehicle (DMSO) was added to the rehydrated diet from 2 days prior to the first intranasal dosing with *Alternaria alternata* extract (Stallergenes Greer - Lenoir, NC - USA). Each day, 1 g of dehydrated food containing either 200 μg of Compound B in DMSO or an equivalent volume of DMSO was provided per 4 g mouse weight. This would correspond to a daily dose of approximately 40 mg/kg, based on the expected food consumption of C57Bl6/J mice [31]. Mice were given an intranasal dose of 20 μg of *Alternaria alternata* extract in 25 μl of PBS, or 25 μl PBS only under general anaesthesia using isoflurane every two days (14 day experiment) (**Fig S2A**) or three times a week (Mon-Wed-Fri) (35 day experiment) (**Fig S2B**). Day 1 was defined as the day of the first *Alternaria* administration. Mice were weighed three times per week. At the end of the experiment, mice were killed by intraperitoneal injection of sodium pentobarbital. Blood was collected into Minicollect tubes (Greiner Bio-one) and centrifuged for 5 min at 10,000 *g* and the serum transferred to a fresh tube. Bronchoalveolar lavage fluid (BALF) was isolated by injection of PBS (3 x 400 μl). Lungs were isolated and digested for 40 min with DNase (50 μg/ml; Sigma) and Liberase (10 μg/ml; Sigma) in RPMI-1640 media, followed by mechanical dissociation through a 40 μm cell strainer, and red blood cells were lysed.

### LPS-induced TNF production

C57Bl6/J mice were fed 1 g of dehydrated food per 4 g of mouse weight containing either 50 μg of Compound B in DMSO or an equivalent volume of DMSO. On day 5, 1 h after the start of the light period, mice received an intraperitoneal injection of either 2 mg/kg LPS (*Escherichia coli* 026:B6, Sigma) in PBS or PBS alone. After 1 h mice were culled by increasing the CO_2_ concentration and confirmed by *per cutane* cardiac puncture. The serum was collected as described above.

### Flow cytometry

For ILC2 experiments, the cell culture supernatant was first collected for cytokine analysis, then ILC2s were diluted 1:5 with PBS containing 1 μg/ml DAPI and Precision Count beads (BioLegend) were added to each sample. Live ILC2 cells for each sample were analysed relative to bead counts using FACSCanto (BD Biosciences). For BALF and lung cell preparations, cell number was determined using the NovoCyte (ACEA Biosciences) and live cells were gated on DAPI negative population. BALF cells were resuspended in 500 μl PBS, 50 μl cell suspension was used for cell counting and the remainder was stained for analysis. One million lung cells were used for analysis. Briefly, cells were pelleted by centrifugation (175 *g* for 5 min) and, following blocking using anti-CD16/CD32, cells were stained with the antibodies in **Table S1**. Data was acquired using FACSCanto and the gating strategy for eosinophils and neutrophils is shown in **Fig S3**. All flow cytometry data was analysed using FlowJo software version 10 (Tree Star).

### Statistical analysis of data

Results are presented as mean and standard deviation. Differences between experimental groups were assessed by statistical analysis as indicated in the figure legends. The results were considered significant if *p* < 0.05; NS not significant, \**p* < 0.05, \*\**p* < 0.01, *** *p* < 0.001. The numbers of mice used in each group for *in vivo* experiments were determined using power calculations based on data from preliminary experiments.

## Results

### A novel SIK inhibitor suppresses the secretion of asthma mediators in mast cells

Compound B is a potent inhibitor of all three SIK isoforms, but is most potent against SIK2. It has improved specificity, compared to HG-9-91-01 and the structurally unrelated MRT199665, compounds that have been used previously to study the roles of SIK family members in mast cells [23].

Compound B suppressed the IL-33-dependent secretion of IL-13 in bone marrow-derived mast cells (BMMC) in a concentration-dependent manner, with complete suppression when it was included in the cell culture medium at 1.0 μM (**Fig 1A**). At this concentration it also suppressed the IL-33-dependent secretion of GM-CSF in BMMC (**Fig 1B**). These results are similar to those obtained using HG-9-91-01 and MRT199665 [23].

**Figure 1.**
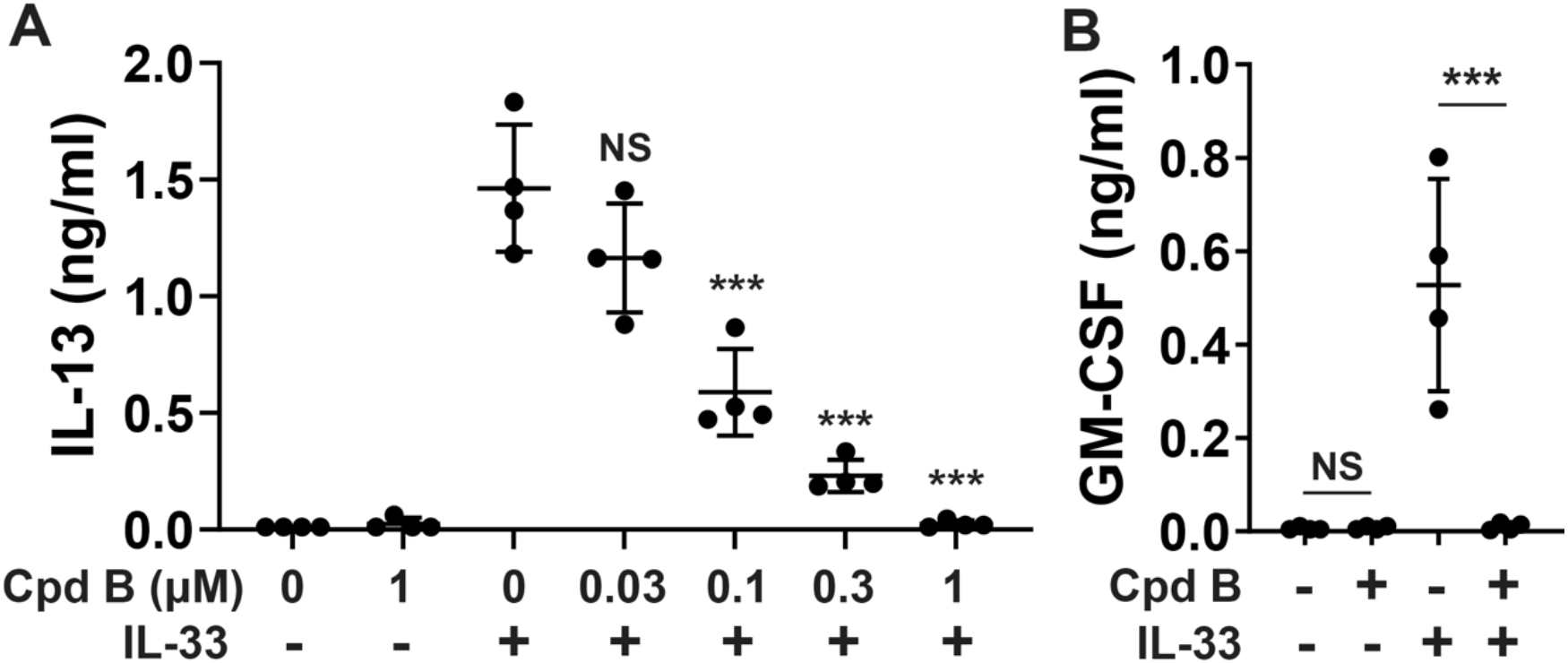
Compound B suppresses the secretion of cytokines from mast cells. BMMC from WT mice were incubated for 1 h without or with Compound B (Cpd B) at the concentrations indicated. **(A)** Cells were stimulated for 2 h with 10 ng/ml IL-33 and the concentration of IL-13 in the cell culture medium was measured. The results shown are the mean and standard deviation from 4 biological replicates and are representative of two independent experiments. Statistical analysis is represented by one-way ANOVA with Dunnett’s multiple comparison test comparing each group treated with Compound B and IL-33 to cells treated with IL-33 only. **(B)** BMMC were incubated for 1 h without or with 1 μM Compound B and stimulated for 6 h with 10 ng/ml IL-33. The concentration of GM-CSF in the cell culture medium was measured. The results shown are mean and standard deviation from four biological replicates and are representative of three independent experiments. Statistical analysis is represented by two-way ANOVA with Sidak’s post-hoc testing; \*\*\**p* < 0.001. (NS, not significant)

### SIK activity is required for IL-33-induced cytokine production by ILC2s

ILC2s are considered to be a significant source of the cytokines that drive allergic asthma, such as IL-13 and GM-CSF [5, 24], and they also secrete IL-6 [25], which appears to have an active role in promoting Th2 cytokine production, eosinophil recruitment and airway hyperresponsiveness in an ovalbumin-induced model of allergic airway inflammation [32]. It was therefore of interest to compare the effect of Compound B on IL-33-stimulated cytokine secretion from ILC2s with the results obtained in mast cells. We found that Compound B reduced the IL-33-stimulated secretion of IL-13 from ILC2s substantially (**Fig 2A**), while the secretion of GM-CSF and IL-6 was reduced partially (**Figs 2B and 2C**), but Compound B had no effect on the secretion of IL-5 (**Figs 2D**). Incubation with Compound B alongside IL-33 stimulation did not affect cell viability (**Fig 2E)**. Taken together, our results indicate that SIK inhibition suppresses the IL-33-dependent secretion of IL-13 and GM-CSF in two innate immune cell types that contribute to asthma pathology.

**Figure 2.**
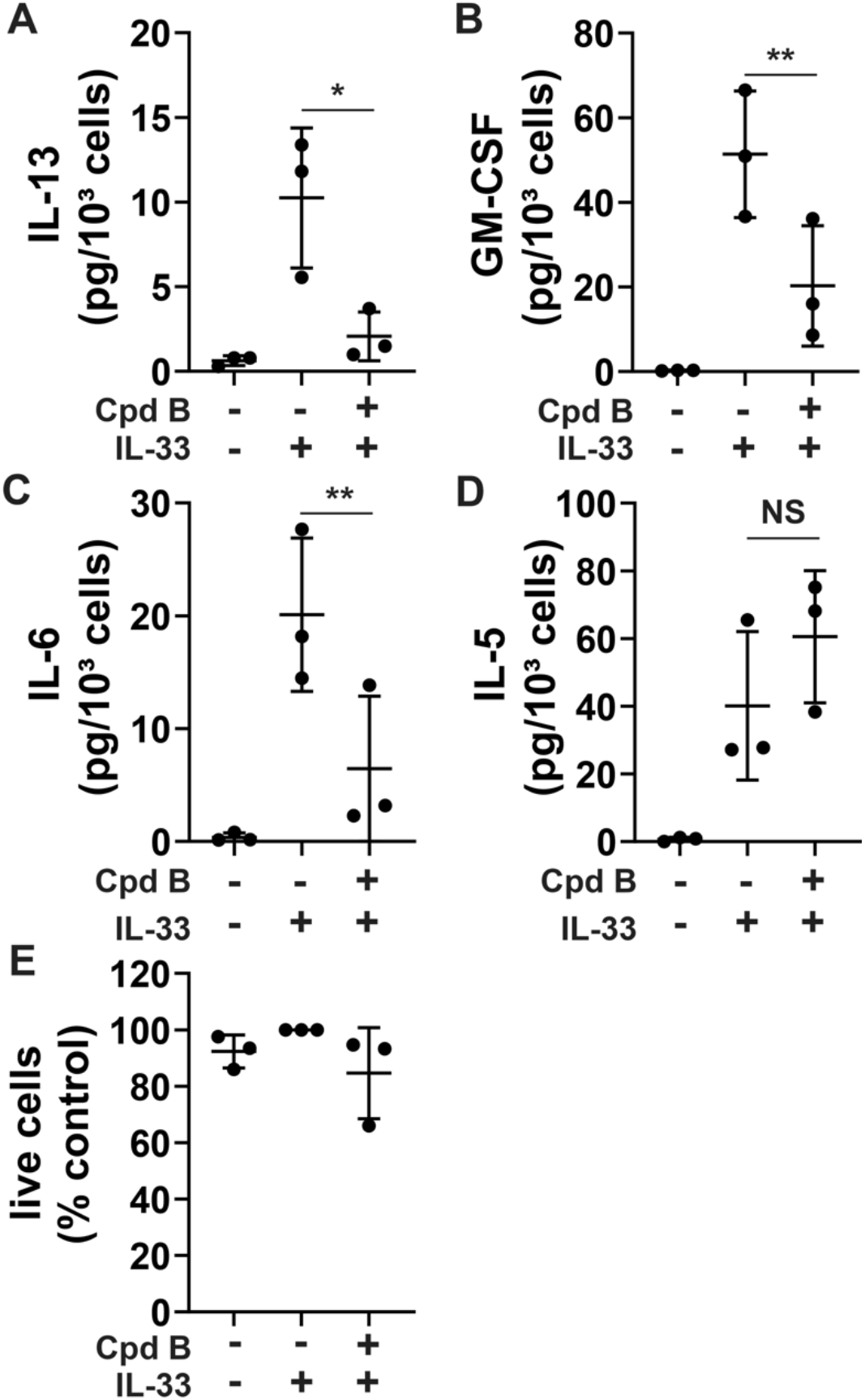
SIK activity is required for maximal secretion of asthma mediators in type 2 innate lymphoid cells. **(A-D)** WT ILC2s were incubated for 1 h without or with 1 μM Compound B (Cpd B) and stimulated for 24 h with 100 ng/ml IL-33. The levels of IL-13 (A), GM-CSF (B), IL-6 (C), and IL-5 (D) secreted into the culture medium were determined. Cytokine secretion for each treatment was adjusted based on number of live cells remaining. **(E)** After removal of the cell culture medium in the experiments shown in A-D, the number of live cells remaining were counted and normalised to IL-33 stimulation in the absence of inhibitors. Cell counts and cytokine secretion were measured in 3-4 technical replicates and the results were averaged. Bars indicate mean and standard deviation and circles show measurements from three independent experiments. The statistics shown represent the results of paired two-tailed t-test comparing IL-33 stimulation in the presence and absence of Compound B; * *p* < 0.05, ** *p* < 0.01. (NS, not significant)

### Compound B suppresses LPS-stimulated TNF production *in vivo*

Compound B has greatly improved pharmacokinetic properties compared to HG-9-91-01 and MRT199665. To check its suitability for *in vivo* studies we studied its effect on LPS-stimulated TNF production in mice. We found that Compound B prevented the LPS-induced production of TNF *in vivo* (**Fig S4**).

### Compound B protects mice in an *Alternaria alternata*-induced asthma model

To test the therapeutic potential of Compound B in a mouse model of allergic lung inflammation, we administered it to WT mice prior to treatment with *Alternaria alternata*. To determine the dose of Compound B required, mice were given the drug after incorporating it into their food to give a predicted dose of 20, 30 or 40 mg/kg per day. Two days after starting compound administration, mice were put on a 14-day *Alternaria* protocol. As expected, eosinophil recruitment into the BALF and serum IgE levels were increased in mice receiving *Alternaria* relative to mice receiving PBS instead of *Alternaria* (**Fig 3**). At doses of 20 and 30 mg/kg, Compound B did not have a significant effect on eosinophil numbers in the BALF or IgE levels in the serum, but at 40 mg/kg a modest reduction in eosinophil recruitment (**Fig 3A**) and a marked reduction in serum IgE levels were observed (**Fig 3B**). A dose of 40 mg/kg was therefore selected for further experiments.

**Figure 3.**
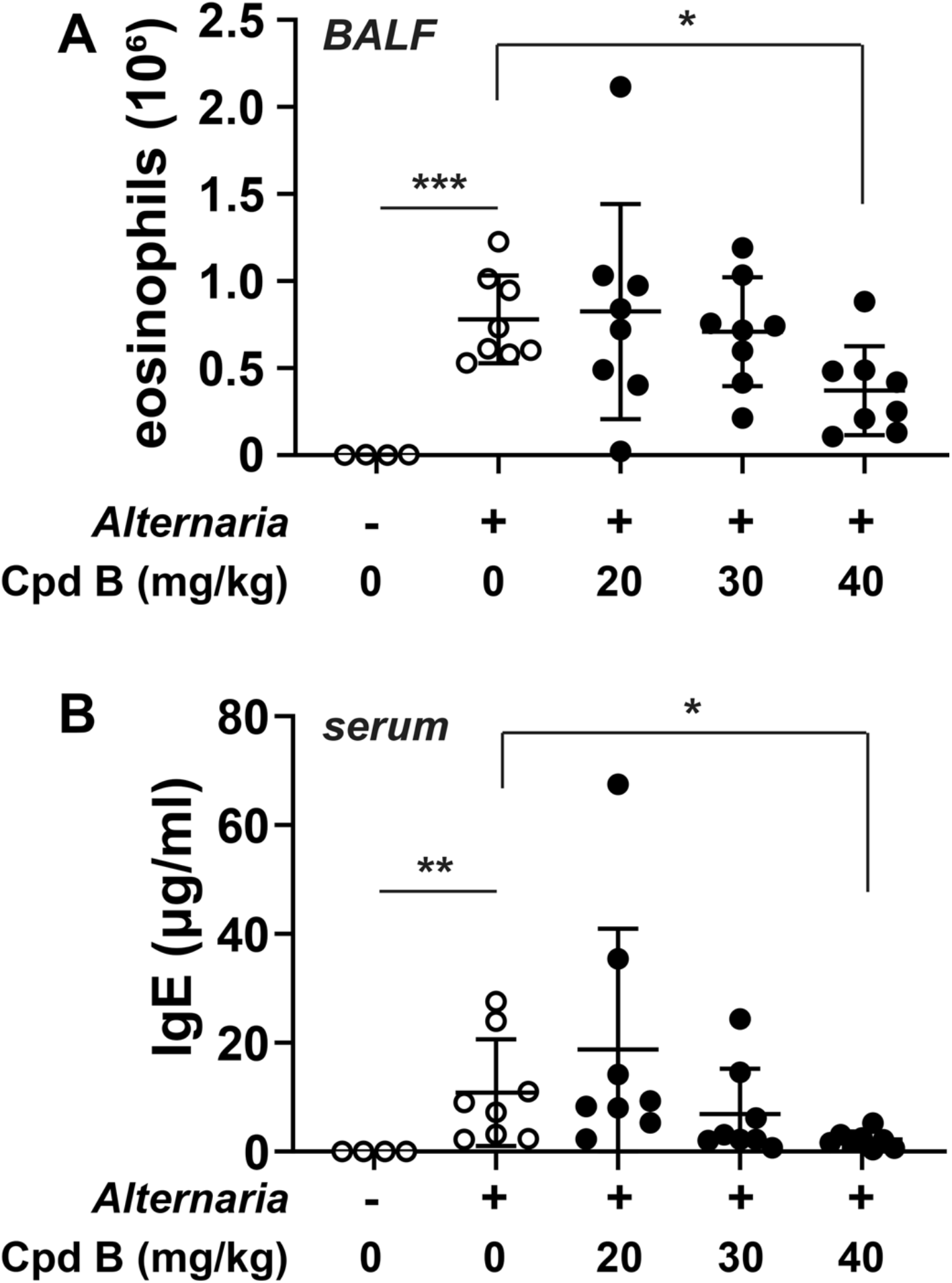
Dose dependent protection against allergic asthma using Compound B. **(A,B)** Mice received 20, 30 or 40 mg/kg of Compound B (Cpd B) or no Cpd B in their diet for 2 days before the start of *Alternaria* administration (see Methods). On day 14, eosinophil numbers in the BALF (A) and serum IgE levels (B) were determined. Bars indicate mean and standard deviation, and circles show measurements for individual mice. Eight mice were used for each *Alternaria* group and four mice were used for the no *Alternaria* control group. Statistical analysis in (A) is represented by Brown-Forsythe and Welch ANOVA test with Dunnett’s T3 multiple comparison test comparing each group to the *Alternaria* group in the absence of Compound B. Statistical analysis in (B) is represented by Kruskal-Wallis test with two-stage linear step-up procedure of Benjamini, Krieger and Yekutieli comparing each group to the *Alternaria* group in the absence of Compound B; \**p* < 0.05, \*\*\**p* < 0.001.

The effects of Compound B were next determined at days 6, 10 and 14 (**Fig S2A**), as well as in a separate 35-day study (**Fig S2B**). *Alternaria* treatment increased the recruitment of eosinophils in both the BALF and lung tissue, which were reduced by Compound B after 14 days (**Figs 4A and 4B**), with more marked reductions observed after 35 days in a separate experiment (**Figs 4C and 4D**). Interestingly, Compound B induced a more rapid increase in neutrophil numbers in the BALF and the lung tissue, which was seen after 6 days following *Alternaria* treatment and maintained up to 14 days (**Fig S5**). Serum levels of IgE were not increased at day 6 following *Alternaria* treatment, but a small increase was seen by day 10 (**Fig 4E**). Larger increases were observed at day 14 (**Fig 4F**) and 35 (**Fig 4G**). Compound B reduced serum IgE levels substantially at both day 14 (**Fig 4F**) and day 35 (**Fig 4G**).

**Figure 4.**
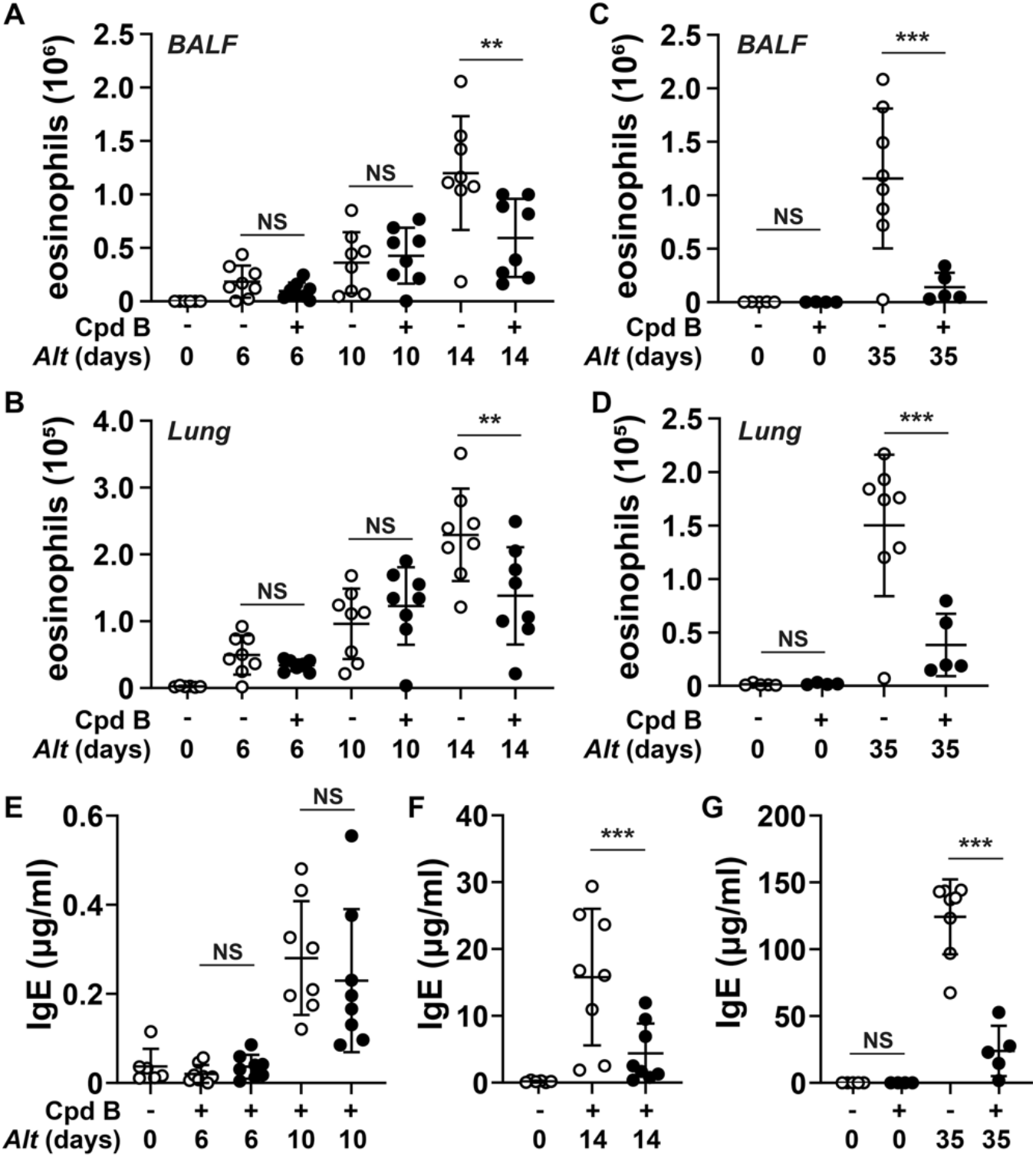
Administration of Compound B protects mice against *Alternaria alternata*-induced allergic lung inflammation. Mice received diet supplemented with either Compound B (Cpd B, 40 mg/kg) or no compound and allergic lung inflammation was induced by repeated dosing with *Alternaria* extract (*Alt*). Two separate experiments were performed, a time course of 6, 10 and 14 days and a separate 35-day experiment. Mice were culled on the days indicated. **(A,B)** Eosinophil numbers in the BALF (A) or lungs (B) following up to 14 days *Alternaria* (*Alt*) administration in the presence or absence of Cpd B treatment. Eight mice were used in each of the *Alternaria* groups and six mice were used in the no *Alternaria*, no Cpd B control group. **(C,D)** As in A,B except eosinophil numbers were determined following 35 days *Alternaria* administration. Data shown is from five mice (no *Alternaria*, no Compound B), four mice (Compound B only), eight mice (*Alternaria* alone) and five mice (*Alternaria* and Compound B). **(E-G)** IgE levels were measured in serum collected from mice following (E) 6 days or 10 days, (F) 14 days or (G) 35 days *Alternaria* administration. Horizontal bars indicate mean and standard deviation and circles show measurements for individual mice. Statistical analysis is represented by two-way ANOVA followed by Sidak’s post-hoc test; ** *p* < 0.01, *** *p* < 0.001. (NS, not significant)

No mortality was observed in any of the mice in the studies lasting up to 14 days. However, in the 35-day study, three mice given both *Alternaria* and Compound B showed acute weight loss, which reached the severity limits permitted in the protocol, requiring their sacrifice before the end of the experiment at day 10 (one mouse) and day 24 (two mice) (**Fig S2C**). The other five mice receiving the compound did not lose weight, and indeed there was an increase in the weight of the animals receiving Compound B (**Fig S2D**). Dosing of Compound B at 40 mg/kg in the food led to 787 nM serum compound levels taken at termination on day 35, 3 h after the final administration of compound in food (**Fig S2E)** which compared to serum Cmax level of 1250 nM in mice dosed by gavage at 10 mg/kg as shown in **Fig S1**.

## Discussion

IL-13 and GM-CSF are thought to contribute to the pathogenesis of asthma (see Introduction) and antibodies neutralising these and other cytokines have been developed and tested in clinical trials as potential treatments for the 5 % of patients with severe asthma who are refractory to the commonly used corticosteroid therapies. However, although these drugs may benefit small subsets of patients, the efficacy of antibodies targeting a single cytokine has proved disappointing in advanced clinical trials for asthma and several such trials have been stopped [33–36]. These findings suggest that either these antibodies need to be trialled in combination, or that a novel drug that suppresses the production of several cytokines that are important contributors to asthma pathology needs to be developed.

In the present study, we used a new SIK inhibitor (Compound B) with improved selectivity and pharmacokinetic properties to show that SIK inhibition reduced the secretion of IL-13 in ILC2s, as well as the secretion of IL-6 and GM-CSF (**Fig 2**). As expected from our earlier studies with other SIK inhibitors [23], Compound B suppressed the IL-33-dependent production of IL-13 and GM-CSF from mast cells (**Fig 1**).

Compound B has improved pharmacokinetic properties compared to other SIK inhibitors and is suitable for *in vivo* studies. We showed that it reduced both eosinophil recruitment and serum IgE levels in an *Alternaria*-induced model of allergic lung inflammation, which are hallmarks of allergic asthma. Interestingly, although Compound B reduced eosinophil recruitment *in vivo*, it did not affect the production of IL-5 in cultured ILC2s, which is thought to be a major chemokine for eosinophil recruitment [37]. Thus, SIKs do not suppress the recruitment of eosinophils to the lung by blocking the secretion of IL-5 from ILC2s. We have shown previously that SIK inhibition reduced CCL24 (eotaxin-2) secretion in mast cells [23], which has been implicated as a chemoattractant for eosinophils in an ovalbumin-induced allergic asthma model [37, 38].

In our earlier studies in mast cells [23], SIK3 was identified as the most important of the three SIK isoforms in controlling IL-13 secretion, although SIK2 activity also contributed to IL-13 production. In addition, SIK3 and SIK2 were critical for the secretion of GM-CSF, TNF and the chemokines CCL2, CCL3, CCL4 and CCL24 in mast cells [23]. In the future, it will therefore be interesting to investigate the therapeutic potential of SIK3-specific inhibitors and dual inhibitors of both SIK2 and SIK3 (that do not affect SIK1) in the treatment of asthma.

It has been reported that prostaglandin E2 (PGE2) signalling via PGE2 receptor 2 has a protective function in allergen-induced asthma (Reviewed in [39]). These effects of PGE2 may be mediated, at least in part, via the SIKs, since the PGE2-PGE2 receptor 2 axis elevates intracellular cyclic AMP and activates cyclic AMP-dependent protein kinase [40], one of whose functions is to phosphorylate and inactivate the SIKs [41, 42].

## Acknowledgements

This study was supported by a Programme grant (MR/R021406/1) from the U.K. Medical Research Council (to P.C.). Compound B has been designed and synthesised by the CRUK Beatson Institute Drug Discovery Unit, which is funded by a Programme Award from CRUK (C7932/A17096) and an MRC DFPS award (MR/M025381/1).

## Author Contribution

M.v.G.B., N.J.D., P.C., L.M., H.J.M and J.S.C.A. conceptualization; M.v.G.B., L.M. and N.J.D. investigation; J.S.C.A. methodology; P.C. and J.S.C.A. supervision; N.J.D. and P.C. writing (original draft); J.S.C.A writing (review and editing); P.C. funding acquisition; L.M., M.S., J.C., M.M., A.S., C.M., J.B. and H.J.M. development of Compound B.

## Abbreviations

BALF: bronchoalveolar lavage fluid
BMMC: bone marrow-derived mast cells
CCL: (C-C motif) ligand
GM-CSF: granulocyte-macrophage colony stimulating factor
IgE: immunoglobulin E
IL: interleukin
ILC2: type 2 innate lymphoid cell
PGE2: prostaglandin E2
SIK: salt-inducible kinase
TNF: tumour necrosis factor
WT: wild type

