## supplementary information for "Salt-inducible kinase (SIK) inhibition is protective in a mouse model of asthma"

**Affiliations**
[1]MRC Protein Phosphorylation and Ubiquitylation Unit and [2]Division of Cell Signalling and Immunology, University of Dundee, U.K., [3]Beatson Drug Discovery Unit, Cancer Research UK Beatson Institute, Glasgow, U.K.

**Supplementary information**

**Table S1** Staining panel for BALF and lung cells.

| **Antigen** | **Conjugation** | **Clone** | **Company** | **Dilution** |
|---|---|---|---|---|
| CD11b | FITC | M1/70 | BioLegend | 1/400 |
| CD11c | APC | N418 | BioLegend | 1/400 |
| CD170 | PE | 1RNM44N | Invitrogen - eBioscience | 1/100 |
| CD19 | APC/Cy7 | 6D5 | BioLegend | 1/200 |
| CD45 | BV510 | 30-F11 | BioLegend | 1/200 |
| CD8 | PE/Cy7 | 53-6.7 | BioLegend | 1/100 |
| Gr1 | BV421 | RB6-8C5 | BioLegend | 1/400 |
| TCRβ | PE/Cy5.5 | H57-597 | BioLegend | 1/200 |

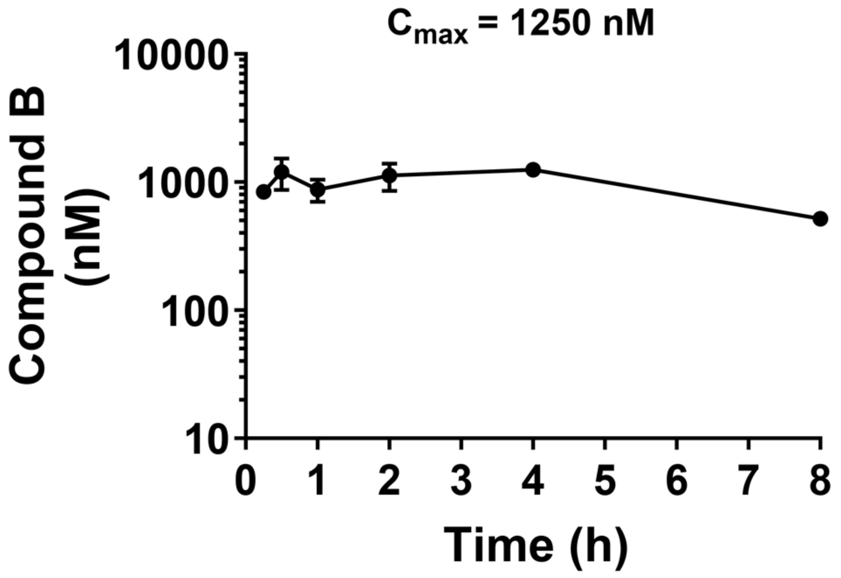


**Figure S1. Pharmacokinetics of Compound B dosed to mice at 10 mg/kg orally by gavage.** A dosing solution of 5 mg/ml Compound B was prepared in 25% propylene glycol/1% Tween/PBS vehicle and administered to CD-1 female mice at 10 mg/kg orally by gavage. Three mice per timepoint were culled at 15 min, 30 min, 1 h, 2 h, 4 h and 8 h after dosing and blood collected for analysis of compound levels. Compound B was shown to have a reasonably flat pharmacokinetic profile over the 8 h study with a $C_{max}$ of 1280 nM at 4 h after dosing.

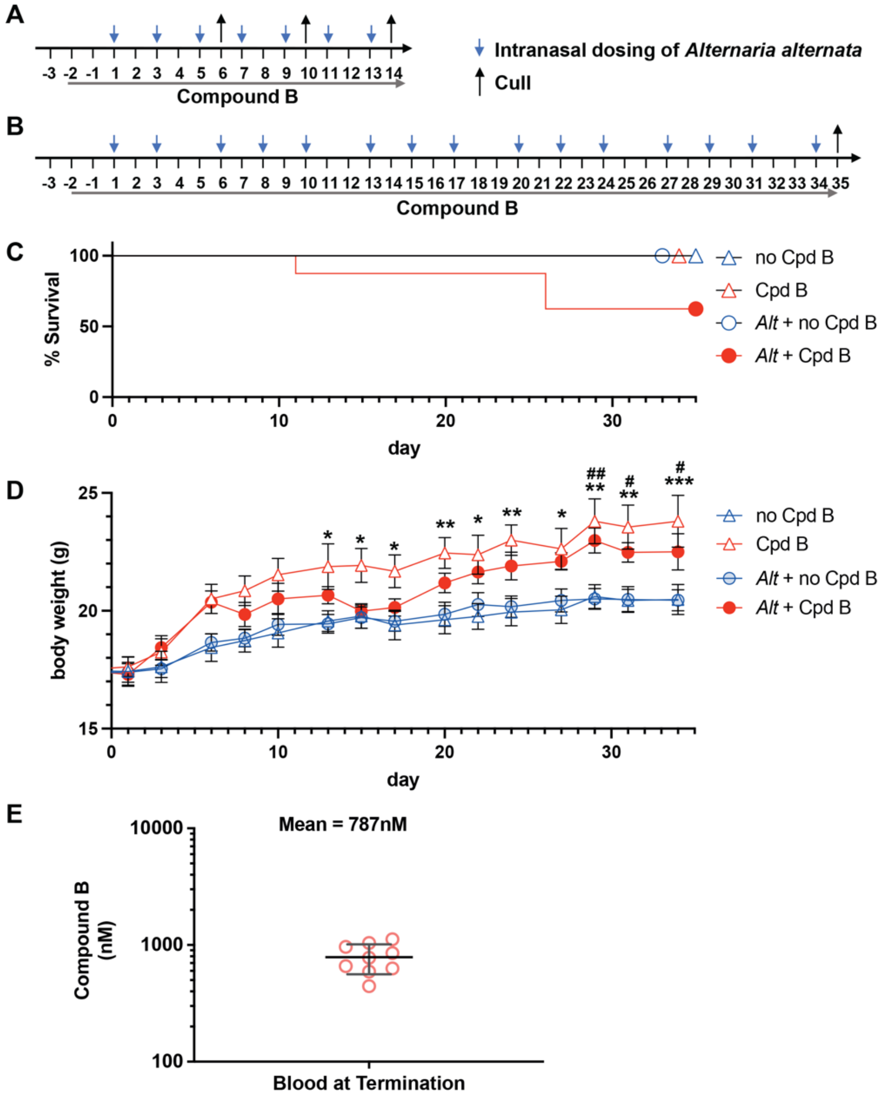


**Figure S2. Schematic showing how Compound B and *Alternaria alternata* were administered in short (14 day) and long (35 day) models.**
Mice received diet supplemented with either Compound B (Cpd B, 40 mg/kg) or no Cpd B and allergic lung inflammation was induced by repeated dosing with *Alternaria* extract (*Alt*) (see Methods). Two separate experiments were performed as shown in the schematic, **(A)** a time course of 6, 10 and 14 days or **(B)** a separate 35-day experiment. Mice were culled on the days indicated. **(C)** Survival and **(D)** weight measurements are shown for the 35-day experiment. In (C,D) eight mice were used in each of the *Alternaria* groups, four mice were administered Compound B only, and five mice were used in the no *Alternaria*, no Compound B control group. Three mice in the *Alternaria* and Compound B group were culled before the end of the experiment due to acute weight loss, these mice are not included in (D). In (D) the data shown are mean and S.E.M. Statistical analysis is represented by two-way ANOVA followed by Tukey's post-hoc testing comparing the effect of Compound B treatment in the absence of *Alternaria* (*) or the presence of *Alternaria* (#). */# $p < 0.05$, **/## $p < 0.01$, *** $p < 0.001$. There was no significant difference between no *Alternaria* and *Alternaria* treatments. (E) On day 35, mice were culled 3h after the final provision on food containing Compound B and serum taken for analysis of Compound B levels. The graph show mean and standard deviation of Compound B concentrations in the 5 surviving mice that received Compound B and *Alternaria* and the 4 mice that received Compound B alone.

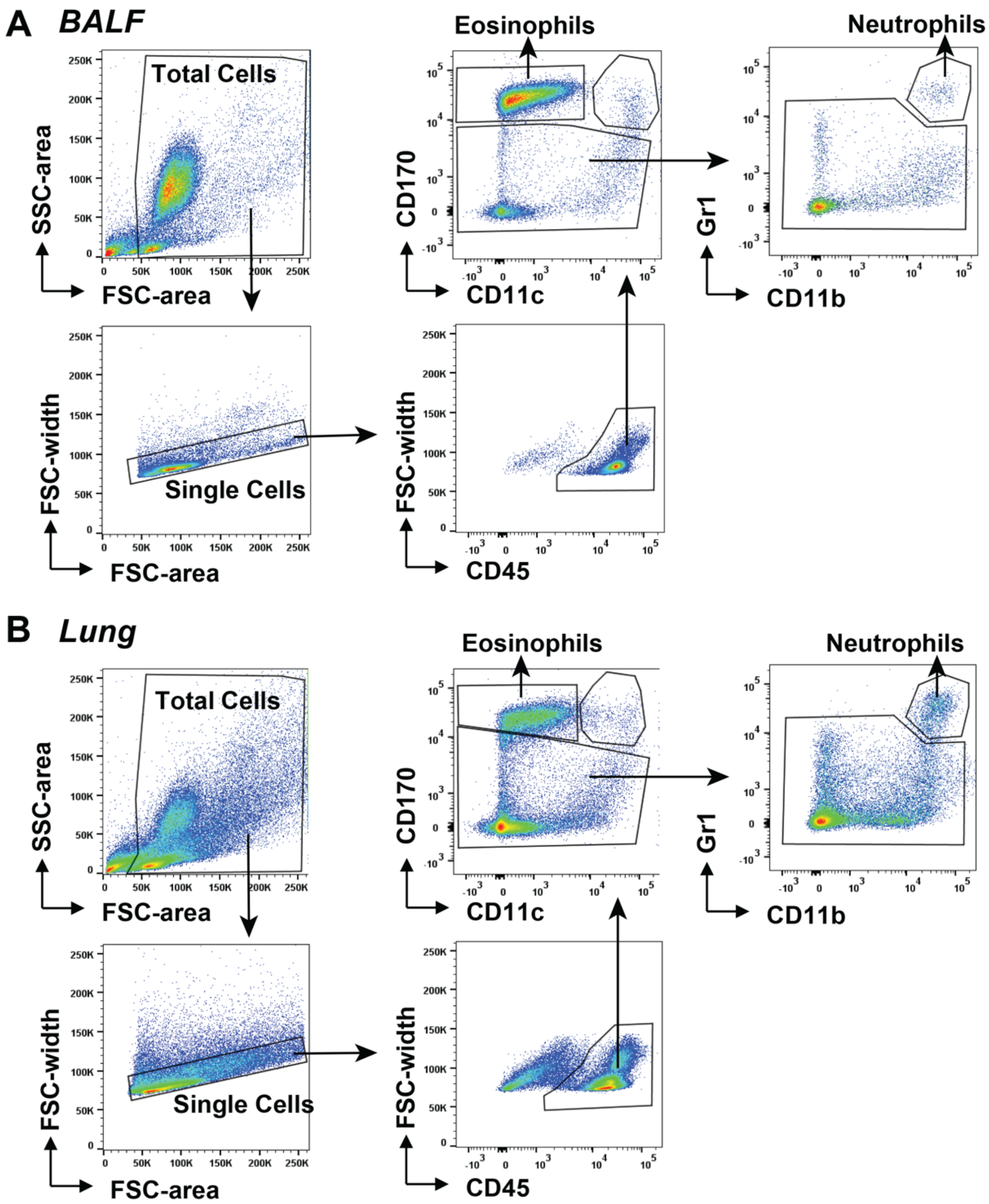


**Figure S3. Gating strategy for flow cytometry of BALF and lung cells.**
BALF cells and lung cells were isolated from mice (see Methods) and analysed by flow cytometry. Cells were pelleted and, after blocking with anti-CD16/CD32, were stained with BV510-CD45, BV421-Gr1, FITC-CD11b, APC-CD11c, PE-CD170, PE/Cy5.5-TCRβ, PE/Cy7-CD8, and APC/Cy7-CD19 antibodies. The gating strategies for BALF **(A)** and lung digests **(B)** are shown. Eosinophils were identified as CD45$^{+ve}$/CD170$^{+ve}$/CD11c$^{-ve}$ and neutrophils as CD45$^{+ve}$/CD170$^{-ve}$/Gr1$^{+ve}$/CD11b$^{+ve}$ cells.

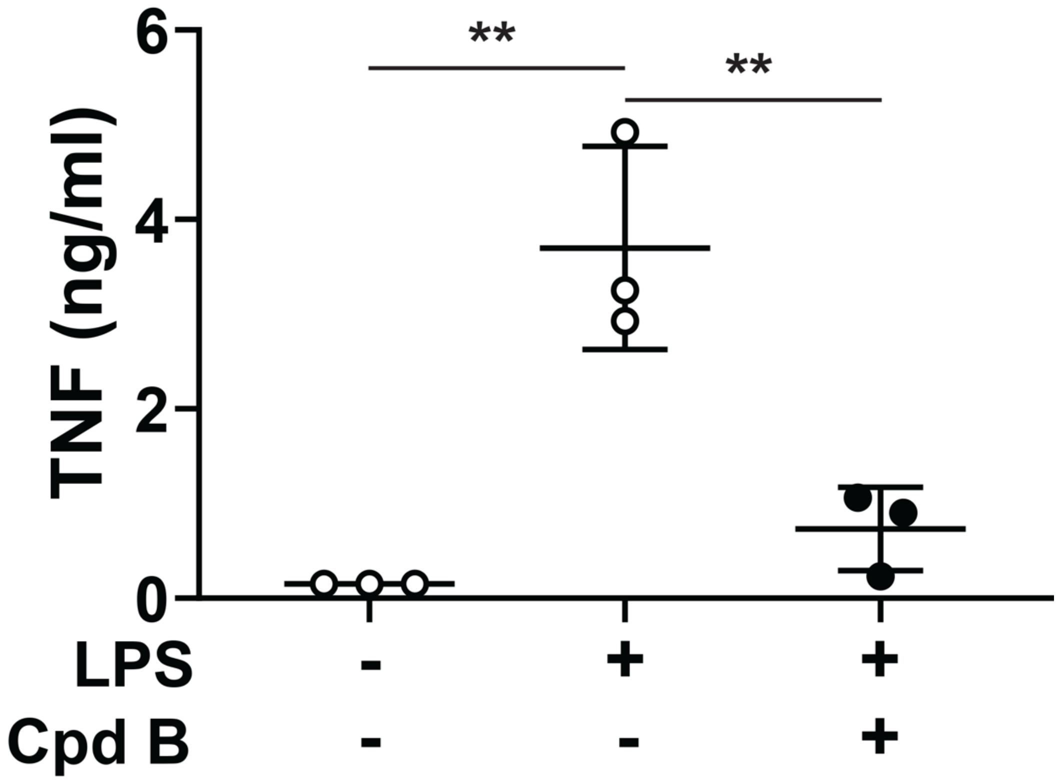


**Figure S4. Compound B blocks LPS-induced TNF production *in vivo*.**
Mice received food containing Compound B (Cpd B, 40 mg/kg) or no drug for 4 days. On day 5, mice received an intraperitoneal injection of 2 mg/kg LPS in PBS, or PBS alone. One hour after injection, TNF levels were measured in the serum. The horizontal bars show mean TNF levels and circles show the TNF levels in individual mice. Three mice were used in each group. Statistical analysis is represented by one-way ANOVA with Tukey HSD post-hoc testing comparing all groups to mice administered with LPS only; ** $p < 0.01$.

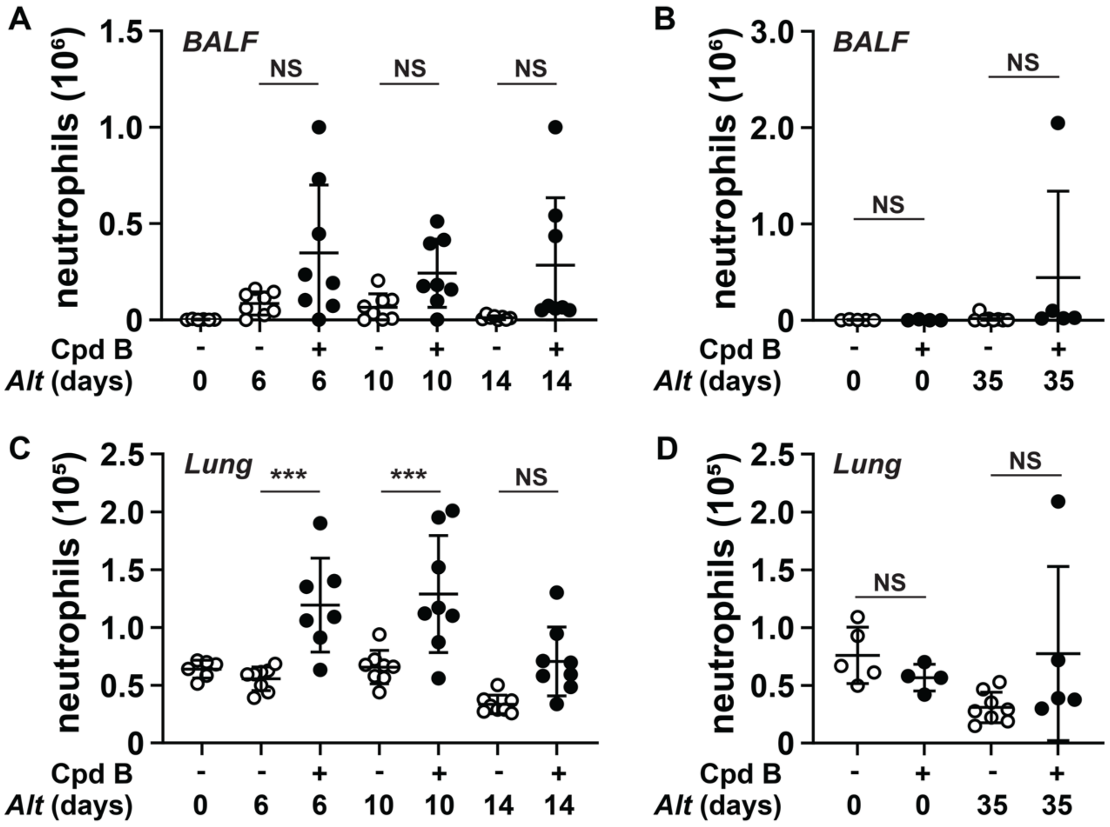


**Figure S5. The administration of Compound B increases neutrophil recruitment following *Alternaria alternata*-induced allergic lung inflammation.**
Data were collected in parallel with those shown in **Fig 4**. Neutrophil numbers in the BALF following **(A)** up to 14 days *Alternaria* (*Alt*) administration or **(B)** 35 days *Alternaria* administration in the presence or absence of Compound B (Cpd B) treatment. **(C,D)** As in A,B except neutrophil number per million cells in the lung was measured. Horizontal bars indicate mean and standard deviation and circles show measurements for individual mice. Group sizes in each experiment are detailed in **Fig 4**. Statistical analysis is represented by two-way ANOVA followed by Sidak's post-hoc testing; *** $p < 0.001$. (NS, not significant)
